# Response diversity inconsistently predicts functional robustness to simulated zooplankton extinctions

**DOI:** 10.64898/2026.06.19.733370

**Authors:** Samuel R.P-J. Ross, Hiromichi Suzuki, Jotaro Urabe, Jamie M. Kass

**Author notes:** **Correspondence:** 1) Samuel R.P-J. Ross, Integrative Community Ecology Unit, Okinawa Institute of Science and Technology Graduate University, 1919-1 Tancha, Onna-son, Kunigami-gun, Okinawa, Japan 904-0495. E 2) Jamie M. Kass, Graduate School of Life Sciences, Tohoku University, Sendai, Japan. E.

## Abstract

Ecosystem functioning can be maintained in species-rich communities even under fairly severe perturbations. This is because communities with high richness include variation both in species’ functional roles (*functional diversity*) and in their responses to environmental changes (*response diversity*). Response diversity has been proposed as a key mechanism underpinning the stabilising role of biodiversity in variable environments. However, less understood is the role of response diversity in stabilising communities against perturbations and thus preserving ecosystem function. Here, we employ community data from 76 reservoirs across the broad latitudinal gradient of the Japanese archipelago to show that zooplankton assemblages with higher variability in responses to environmental variables can retain functional diversity as species are removed, but that the result depends on the variable examined. We combine empirically derived occurrence data for 47 zooplankton species with biotic and abiotic variables in a joint species distribution model to derive species-specific environmental responses, then measure response diversity to environmental axes including fish community structure and water temperature. We also measure functional trait diversity of zooplankton assemblages and simulate sequential species extinctions, capturing the extinction thresholds beyond which half the functional diversity is lost. Finally, we combine these data streams to show that response diversity can predict higher functional robustness in zooplankton assemblages, but not consistently. The role of response diversity in predicting functional robustness was contingent on the specific metric and environmental variable considered. We found that a balance of positive and negative species responses to water temperature was a significant predictor of robustness, though other metrics and environmental variables mainly yielded non-significant relationships. Overall, we show that response diversity can confer stability to perturbations such as species extinctions, and we demonstrate the utility of species distribution models for measuring response diversity, overcoming mechanistic data limitations and expanding the toolkit available for studying response diversity in natural systems.

## 2 Introduction

Global environmental change presents a major challenge to the continued delivery of ecosystem functions and services. Understanding the drivers of ecosystem stability amid environmental change is therefore a central goal of contemporary ecology. The insurance effect posits that biodiversity both enhances and stabilises ecosystem functioning in variable environments (Yachi and Loreau 1999; Loreau et al. 2021). This stabilising effect occurs through a variety of potential pathways, including compensatory or asynchronous population dynamics among species (Shoemaker et al. 2022) and response diversity, or the range of species’ responses to environmental change (Elmqvist et al. 2003; Mori et al. 2013). Response diversity is high when species respond differently to disturbance and low when they respond similarly. Ecosystem functions of communities with higher response diversity should be less variable under directional environmental change because declines of sensitive species can be offset by tolerant species or disturbance specialists (Yachi and Loreau 1999; Mori et al. 2013). In this way, response diversity may capture how biodiversity confers stability in variable environments.

The multiple dimensions of ecological stability—including, for example, axes of temporal variability, and resistance to and recovery from perturbations (Donohue et al. 2013)— necessitate a holistic understanding of how response diversity contributes to stability across these dimensions (Ross and Sasaki 2024). Given the focus of biodiversity insurance theory on the variability of ecosystem processes and functions through time (Yachi and Loreau 1999), most studies of response diversity to date have focused on its relationship with temporal variability. Higher response diversity has been linked to temporal stability for diverse taxa such as grassland plants (Sasaki, Lu, et al. 2019), trees (Schnabel et al. 2021), birds (White et al. 2023), and aquatic protists (Leary and Petchey 2009). However, response diversity has also been conceptualised as a driver of ecosystem resilience (Elmqvist et al. 2003; Mori et al. 2013) and related stability dimensions including resistance and robustness (Ross and Sasaki 2024; Ross, Barros, et al. 2026).

Mori et al. (2013) showed that the relationship between species’ environmental responses and their functional roles underpins how response diversity can drive resilience. By building on the response-effect trait framework, which classifies species’ functional traits as related to functioning and/or environmental responses (Suding et al. 2008), De Bello et al. (2021) conceptualised how correlations between species’ responses and their functional effect traits shape different dimensions of stability, such as robustness. Functional robustness is the ability of a community to maintain ecosystem functioning in the face of extinctions, and it can be estimated by measuring the rate at which functional diversity declines as species are lost from the community (Petchey and Gaston 2002a; Ross, Arnoldi, et al. 2021). Owing to sampling effects, greater richness increases the likelihood of the presence of species that have both diverse functional roles (high functional diversity) and varied environmental tolerances and responses (high response diversity). As a result, response diversity and functional diversity are expected to be positively correlated (Mori et al. 2013). In such cases, the net effect of environmental perturbations on communities should, on average, be smaller in species-rich communities because environmental tolerances are more varied and the loss of functional diversity is less severe (functional robustness is higher). Thus, when response diversity and functional diversity positively covary, response diversity provides a mechanistic explanation for how species richness buffers ecosystems against losses of functional diversity (Elmqvist et al. 2003; Mori et al. 2013; De Bello et al. 2021). However, given the scarcity of studies comparing the two measures of diversity, we still lack a comprehensive understanding of their relationships to each other and to functional robustness in empirical systems.

Response diversity studies tend to either quantify response diversity based on species’ functional response traits (Sasaki, Lu, et al. 2019; White et al. 2023), which are assumed to relate to environmental responses (Suding et al. 2008; Oliver et al. 2015), or derive environmental responses from mechanistic experiments of species’ functional responses (Leary and Petchey 2009; Polazzo, Hämmig, et al. 2025). The decision of whether to use direct or indirect measures of environmental responses is often dictated by study design. Although direct measures are ideal for dissecting functional responses, experimentally deriving environmental responses for many species or measuring many functional traits in the field may be impractical, particularly when studies require large sample sizes and replication. In this light, how might we measure response diversity at landscape scales? Genung and Winfree (2025) and other studies identify response diversity based on a modelled interaction term between species and agricultural land cover, suggesting response diversity occurs if species’ slopes differ significantly (see also Winfree and Kremen 2009; Cariveau et al. 2013). Another option may be to predict species’ environmental responses over time and space using niche models trained on occurrence records and environmental predictors (e.g., Antão et al. (2022)). As these are correlative methods, they do not include the physiological mechanisms underpinning environmental responses, nor can they make mechanistic forecasts of the effects of environmental change on species’ functional contributions. However, these methods are much more scalable over space and time and can provide valuable information about the role of biotic and abiotic environmental responses in structuring ecological communities (through species occurrences if not functional responses). Here we indirectly infer species’ environmental responses based on how occurrences scale with a set of empirically-derived environmental variables to estimate response diversity within the bounds of our methodological toolkit (Ross, Barros, et al. 2026).

Here, we use extensive zooplankton assemblage survey data from 76 reservoirs (human-made lakes) across the broad latitudinal gradient of the Japanese archipelago to test the hypothesis that zooplankton assemblages with higher response diversity are more functionally robust to species extinctions (Figure 1). We calculate response diversity based on estimates of species’ environmental responses from a joint species distribution model, and functional diversity using functional trait data for each assemblage. We then measure functional robustness by simulating removals of species from the assemblage and finding the proportion of extinctions required to cause a 50% decline in functional diversity. Species extinctions represent a major axis of human impacts on ecosystems (Diamond 1989), and high functional robustness is required for communities to maintain ecosystem functioning amid the ongoing biodiversity crisis (Petchey and Gaston 2002a; Mumby et al. 2014). If species’ functions and responses are correlated, we expect ecosystem functioning for assemblages with high response diversity to be more robust to simulated species extinctions.

**Figure 1:**
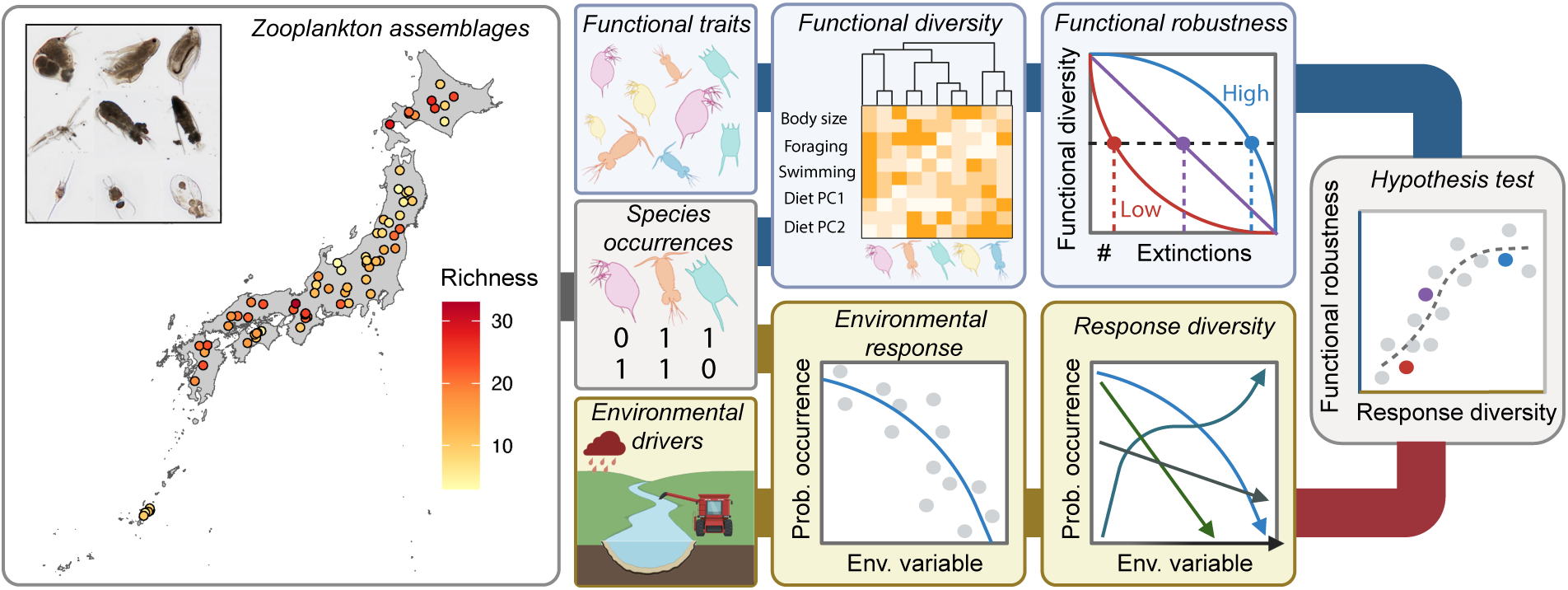
Workflow diagram for measuring response diversity and functional robustness. We join species occurrence records for zooplankton from 76 dammed reservoirs across the Japanese archipelago with functional trait data and environmental drivers. Using functional trait data for species present per site, we measure functional diversity and its robustness to species extinctions. Using joint species distribution models of zooplankton species, we predict species’ environmental response curves, then estimate the diversity of these responses among species within a site. Finally, we test the relationship between response diversity and functional robustness. Created with BioRender.

## 3 Methods

All analyses were performed with the R programming language v4.5.1 (R Core Team 2025).

### 3.1 Zooplankton assemblages

We examined zooplankton assemblages in reservoirs across a broad latitudinal gradient in the Japanese archipelago (longitude: 127.952–143.387, latitude: 26.480–44.114) spanning a wide elevational range (28–976 m; Figure 1). Zooplankton data were routinely collected in 96 reservoirs by the National Census on River and Dam Environment (5th round) conducted from 2011 to 2015 by the Ministry of Land, Infrastructure, Transport and Tourism of Japan. These data are available via the River Environmental Database (https://www.nilim.go.jp/lab/fbg/ksnkankyo/index.html). In this census, zooplankton samples were collected at sites near the dam or at the deepest point of the reservoir using 100-*µ*m mesh conical plankton nets and bottle samplers such as Van Dorm sampler employed from the bottom or mid-depth to the surface. Since conical plankton nets capture a wider range of taxa than bottle samplers (Suzuki, Osugi, et al. 2026), we used zooplankton community data collected via net sampling. We confirmed taxonomic classifications based on Suzuki, Osugi, et al. (2026). As sampling volume was different between the reservoirs, we did not consider abundance data for zooplankton taxa, and instead reduced counts to binary presence/absence. Then, for each reservoir, we combined the time-series data into a single presence/absence value per species. To avoid the influence of rare taxa on the overall assemblage structure, we filtered out zooplankton species present in fewer than 10% of all reservoirs (n = 8). After removing sites with missing abiotic and biotic environmental variables (see below), we were left with 76 reservoir sites for analysis.

### 3.2 Environmental variables

We assembled a range of environmental variables representing reservoir characteristics, elevation, nutrient availability, water temperature, and fish community structure (Figure S1). For reservoir characteristics, we obtained watershed area, effective storage volume, dam height, and dam width from the Japan Dam Foundation database (http://damnet.or.jp/ Dambinran/binran/TopIndex.html) and periodic reports issued by each reservoir management office. We acquired reservoir elevation data from Japan’s Database of Dams (http://mudam.nilim.go.jp/home). Finally, we obtained surface water temperature (*^o^C*) and nutrient data on total phosphorus (mg/L), total nitrogen (mg/L), and chlorophyll-*a* concentration (*µ*g/L) per reservoir from Japan’s River Environmental Database (http://www.nilim.go.jp/lab/fbg/ksnkankyo/index.html). We summarized these abiotic variables by taking the mean annual value for the years in which zooplankton were collected. To reduce collinearity and the number of predictor variables for modelling, we ran separate principal components analyses (PCAs) for the reservoir variables (Table S3) and nutrient variables (Table S4), keeping the first axes. The final variable set (reservoir PC1, elevation, nutrient PC1, and water temperature) had relatively low collinearity (Spearman’s *ρ <*0.6).

For biotic variables, we used axes from a community ordination to represent fish community structure. We obtained presence-absence data for fish species documented in each reservoir between 2000 and 2010 from the River Environmental Database. As no data were available for specific years, we combined records to derive single presence-absence values of fish species per reservoir. As with zooplankton species, we filtered out fish species with fewer than 10% presences among all reservoir sites, retaining a total of 35 fish species (Table S5). We then calculated pairwise Sørensen distances for the fish species and ran a principal coordinates analysis (PCoA) with the package vegan v2.7-1 (Oksanen et al. 2025), retaining the first three axes as predictor variables, which explained 40.2% of the variance in the fish community (Table S5). On the first axis (PCoA 1), cyprinid fishes such as *Zacco platypus* and *Micropterus salmoides* exhibited high positive scores, whereas *Oncorhynchus mykiss* and *Salvelinus* sp. showed high negative scores. We therefore interpreted the first axis as reflecting the prevalence of warm-water fish. On PCoA 2, species that prefer flowing water, such as *Leuciscus hakonensis* and *Plecoglossus altivelis*, exhibited high scores; accordingly, this axis was interpreted as representing riverine fish. Conversely, on PCoA 3, species that prefer still water, such as *Misgurnus anguillicaudatus* and *Hypomesus nipponensis*, exhibited high scores, and this axis was therefore interpreted as representing lentic fish communities (see Figure and Table for site and species loadings on the PCoA axes).

### 3.3 Species responses and response diversity

We used a joint species distribution model (JSDM) to estimate the responses of zooplankton species to environmental variables. We employed the Hierarchical Modelling of Species Communities (HMSC) framework, implemented in the package Hmsc v3.0-13 (Tikhonov et al. 2020), to fit Bayesian joint probit regression models to the zooplankton presence-absence data. We compared five models ranging from simple (few variables) to complex (many variables) trained on different variable combinations (Table 1), then selected an optimal model based on performance metrics. We assessed model convergence by inspecting coefficient effect sizes and Gelman and Rubin’s convergence diagnostic (Brooks and Gelman 1998). We compared model performance using WAIC, an information criterion metric used for hierarchical models that penalises models with more predictor variables (Watanabe and Opper 2010). For models with less than two units difference for WAIC, which are treated as statistically similar in performance, we compared results of spatial cross-validation. This tests the ability of the model to transfer to conditions likely outside the training range, thus evaluating the model’s ability to extrapolate (Roberts et al. 2017). We defined four spatial folds using a systematic selection (Supporting Figure S2) with the cv spatial function in the blockCV package v3.1-5 (Valavi et al. 2018). We calculated validation AUC for each cross-validation fold and averaged these values per species. For models with similar WAIC, we compared the averages of species’ AUC values. Before calculating response diversity metrics, we filtered out species with AUC values that were less than 0.6. This value was chosen as a conservative threshold, as values below 0.5 denote performance worse than random.

**Table 1:** Joint species distribution models fit to zooplankton species in reservoirs across Japan. . Variables are abbreviated as follows: T = water surface temperature, R = reservoir variables PC1, E = elevation, N = nutrients PC1, F1-3 = fish community PCOA1-3. Models were evaluated via WAIC, but as all had *<* 2 units difference, they were compared with median validation AUC calculated with spatial cross-validation. Model 5 was chosen as optimal based on median AUC. Before calculating response diversity metrics, we removed species with *<* 0.5 AUC, resulting in a median AUC of 0.71.

| Model | ΔWAIC | median AUC |
| --- | --- | --- |
| 1. $T + T^2$ | 0.48 | 0.55 |
| 2. $T + T^2 + R$ | 0.31 | 0.54 |
| 3. $T + T^2 + R + E$ | 0.07 | 0.54 |
| 4. $T + T^2 + R + E + N$ | 0 | 0.61 |
| 5. $T + T^2 + R + E + N + F1 + F2 + F3$ | 0.94 | 0.66 |

After selecting an optimal model, we used the computeVariancePartitioning function in Hmsc to calculate the relative contribution of different environmental variables to structuring zooplankton occurrence across Japan (Antão et al. 2022), then ranked variables by their frequency as the variable with the highest proportion of variance among all species. We concentrated on the variables with highest overall importance for subsequent response diversity analyses. We then generated marginal response curves per species by predicting probabilities of presence over gradients of each environmental variable (Figure 2A,B), then computed their first derivatives to calculate response diversity metrics.

**Figure 2:**
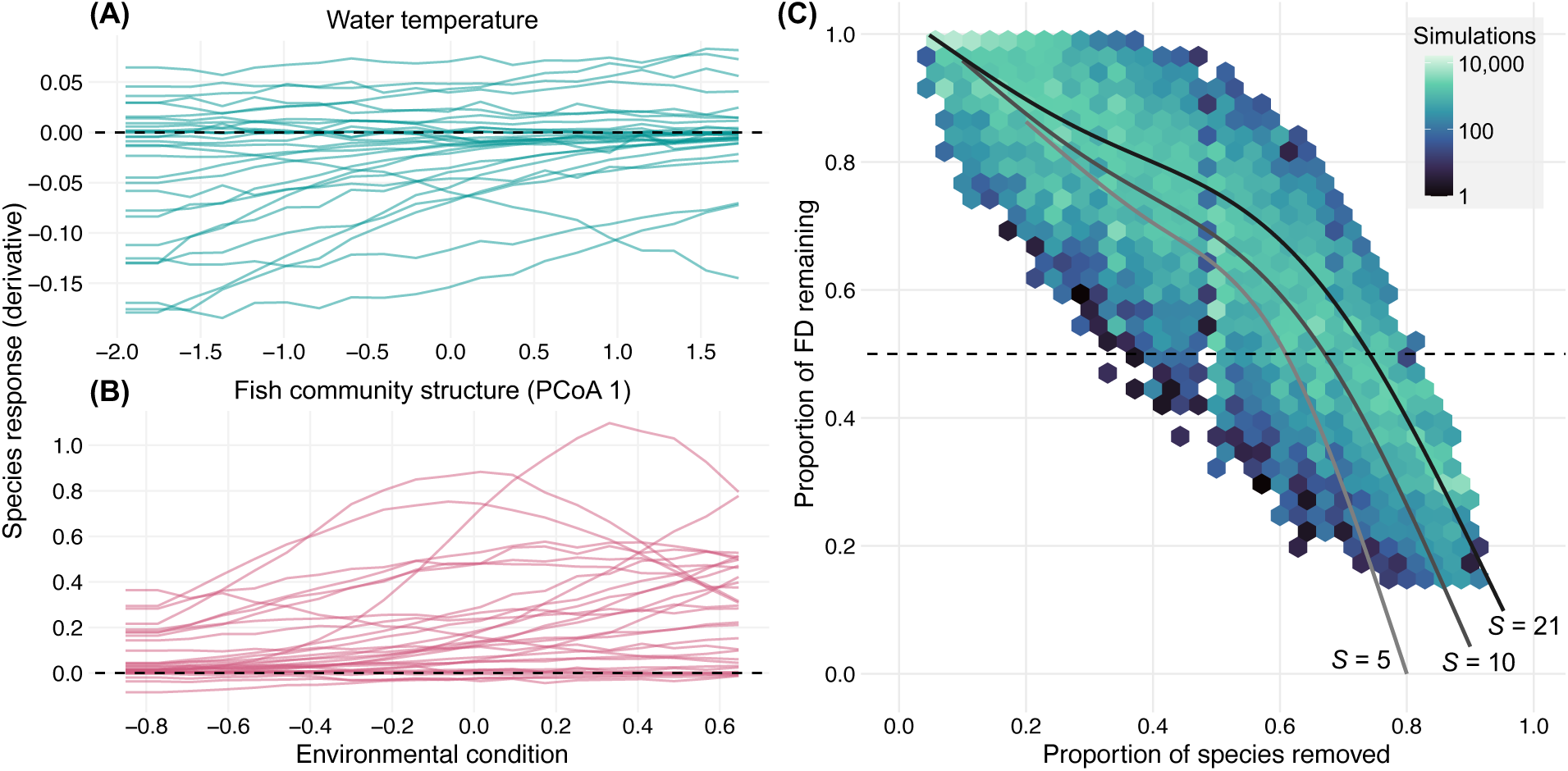
Species-environment relationships and extinction simulations. Species’ responses to (A) water temperature and (B) fish community structure (PCoA 1), calculated from the outputs of joint-Species Distribution Models as the probability of occurrence under a given environmental condition. Individual lines represent different zooplankton species (all species from the regional pool) with response diversity calculated based on the presence/absence of different species per reservoir assemblage. (C) Simulated extinctions on empirical zooplankton assemblages for the calculation of functional robustness (proportion of extinctions required to bring about a 50% loss in functional diversity). Each assemblage underwent 1,000 random extinctions per possible richness level. Hexagonal bins show the density of outcomes across all simulations, while lines indicate smoothed trends (Gaussian process smoother with *K* =4) of functional diversity loss for representative low, medium, and high initial species richness levels (*S* = 5, 10, and 21).

We measured response diversity with three complementary metrics based on the first derivatives of species’ environmental responses (Ross, Petchey, et al. 2023): dissimilarity, diver-gence, and imbalance. Response dissimilarity measures the magnitude of differences among species’ environmental response curves (Ross, Petchey, et al. 2023), equivalent to Leinster and Cobbold (2012)’s similarity-based diversity metric with the sensitivity parameter tuned to species richness only (*i.e.*, ignoring species abundances; *q* = 0). Dissimilarity scales from 1 to *S*, the total species richness: this is lowest (1) when all species respond identically to an environmental variable, and highest (*S*) when slopes of species’ responses are as dissimilar as possible. Response divergence focuses on the direction rather than the magnitude of the response (Ross, Petchey, et al. 2023). Divergence is maximal (1) when species responses are symmetrical around zero (i.e., positive and negative responses balance each other out). Response imbalance is a newly developed metric that measures how imbalanced the sign and magnitude of species’ responses are to environmental changes (Polazzo, Hämmig, et al. 2025). Here, we calculated response imbalance as:

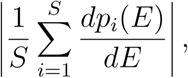

where *p_i_* is the vector of occurrence probabilities of species *i*, *E* is the vector of values of an environmental variable, and *S* is the species richness of the community. Calculated this way, response imbalance measures the balance of instantaneous responses along an environmental gradient (based on derivatives, *d*) instead of fora specific environmental change as in Polazzo, Hämmig, et al. (2025). Values of response imbalance approaching zero are expected to be stabilising owing either to balanced positive and negative environmental responses (species asynchrony) or weak environmental responses (population stability), while high response imbalance is expected to reduce stability (Polazzo, Hämmig, et al. 2025). Finally, we also included mean species responses—the mean of the derivatives of all environmental response curves per assemblage—as an additional predictor of functional robustness, since mean environmental responses are important for stable coexistence under environmental change (De Laender et al. 2023) and temporal stability under pulse disturbances (Kunze et al. 2026). In all cases, we measured instantaneous response diversity for each value of each environmental axis and summarised response diversity as the mean for each environmental variable per reservoir (Ross, Petchey, et al. 2023). The calculations for imbalance and mean response are similar, though as imbalance values are absolute (*i.e.*, positive) values, response imbalance and mean response values differ when communities include both positive and negative environmental responses among species.

### 3.4 Functional traits and robustness

We evaluated the functional diversity of each zooplankton assemblage based on several functional traits from the literature. As zooplankton trait data were not measured in the national census, we determined trait values for the adult stage of each taxon based on several sources (Table S2). We used body size, food preference, feeding strategy, and swimming mode as functional effect traits, since they are known to affect the functional roles of these species rather than their responses to environmental change (Suding et al. 2008; Oliver et al. 2015). Food preference was represented as the first two axes of a principal component analysis (PCA) on five dietary composition variables: blood-feeding, carnivory, detritivory, herbivory, and omnivory (Salt et al. 1978; Suzuki, Ichiyanagi, et al. 2025). Each variable scaled from 0–100, indicating the approximate percentage of a species’ diet following that category; dietary variables were thus highly correlated before the PCA ordination. We scaled continuous traits (body size and the two dietary composition axes) by their standard deviations to prevent differences in relative importance.

We measured functional diversity using Petchey and Gaston (2002b)’s FD metric, a tree-based approach. The FD metric is ideal for studying functional robustness because it cannot increase as species are removed from a community (Petchey and Gaston 2002b) and it has previously been used to quantify declines in functional diversity under simulated extinctions (Petchey and Gaston 2002a). We first measured the Gower distance between all species pairs in multidimensional trait space (Pavoine et al. 2009), then constructed a minimum-spanning functional dendrogram based on these distances, where branch lengths represent the distances between species in trait space (Petchey and Gaston 2002b). Finally, we measured FD for each reservoir site as the sum of the branch lengths of the dendrogram for the species in that reservoir’s assemblage using the *BAT* package 2.11.0 (Cardoso et al. 2025).

For each richness level below the observed richness of each assemblage (*i.e.*, *N* = 1, 2, 3 for richness of 4), we simulated perturbations by randomly subsampling species from the assemblage 1,000 times and recalculating FD for each iteration. This produced 1,000 bootstrapped subassemblages for each site that decline in FD as the perturbation strength increases (and species richness decreases). Across all sites, these simulations produced expected patterns of FD declines with species extinctions (Figure 2C) based on previous work (Petchey and Gaston 2002a). Finally, we measured functional robustness as the proportion of species extinctions required to reduce functional diversity to ≤ 50% of the full assemblage (Domínguez-García et al. 2019; Ross, Arnoldi, et al. 2021).

### 3.5 Data analysis

We used the estimates from above to test our hypothesis that response diversity (RD) promotes functional robustness (FR). First, to establish whether species’ responses and effects are correlated—which informs whether we expect RD to drive FR (De Bello et al. 2021)—we fit ordinary least-squares models between RD and functional diversity (FD) and compared model fit between a linear model and a second-order polynomial model using AIC.

For each RD metric separately, we fit linear models to model relationships among species richness (S), RD, FR, and FD. The models were: *RD ↑ S*, *FD ↑ S*, *FR ↑ RD*+*FD*+*S*. We tested model residuals for spatial autocorrelation with Moran’s I tests using the R package spdep 1.4-1 (Bivand and Wong 2018), and if a significant result was found (*p <* 0.05), we refit the model using generalised least squares with spatial Gaussian correlation structures (longitude and latitude predictors) with the R package nlme 3.1-168 (Pinheiro et al. 2025). We then fit piecewise structural equation models (SEMs) based on these linear models using the R package piecewiseSEM 2.3.0.1 (Lefcheck 2016) to determine the pathways through which richness (S) affects robustness (FR), including via response diversity (RD).

To assess the significance of the observed relationships in the SEM path diagram, we conducted random permutation tests and compared the observed values to null distributions. For each permutation, we first shuffled the values of S, RD, FD, and FR 1,000 times, then refit the models for all variables and re-ran the SEM; these simulations were combined to derive a null distribution for each path coefficient (Suzuki, Miyamoto, et al. 2025). Then, focusing on the relationship between RD and FR, we tested whether the observed value of the path coefficient fell outside the 95% confidence intervals of the null distribution. An observed value outside this range indicates that RD has a significant effect (two-tailed *p <* 0.05) on FR after accounting for the direct effect of S on RD.

## 4 Results

All JSDMs had good convergence, with effective sizes close to the number of posterior samples and Gelman and Rubin’s multivariate convergence statistics close to 1. We selected the JSDM with all eight variables (model 5 in Table 1) based on mean validation AUC (averaged across species), as WAIC values were all within two units. For this model, 16 zooplankton species resulted in AUC values less than 0.6, so we removed these from the subsequent diversity calculations. This resulted in four dams with fewer than three species each, so these were also removed, leaving us with 72 dams total. Variance partitioning showed that two variables had the highest variance proportions among all species: fish community PCoA 1 (highest for 49% of species) and water temperature (highest for 32% of species), so we focus on these two variables for subsequent analyses and report results for the others in the Supporting Information. Species varied considerably in their responses to these environmental variables (Figure 2). Since fish community PCoA 1 represents the prevalence of warm-water fish, most of which are omnivorous and prey on zooplankton (such as for cyprinid fish), a high PCoA 1 score likely implies high predation pressure for the zooplankton community (Jeppesen et al. 1997).

We first tested whether functional (effect) diversity and response diversity were positively related, forming the rationale for testing whether response diversity promotes functional robustness (De Bello et al. 2021). For both our main environmental variables—water temperature and fish community composition—response diversity was significantly related to functional diversity (Figure 3). We found that the imbalance of species’ environmental responses to water temperature was negatively related to functional diversity (slope = *-*22.0 *±* 7.73, *F* = 8.13, *p* = 0.006, *R*^2^ = 0.09; Figure 3A): assemblages with more balanced water temperature responses were typically more functionally diverse. Response imbalance of species’ environmental responses to fish community structure was also related to functional diversity, with a second order polynomial model outperforming a linear model (*F*-test *p >* 0.05), such that medium values of response diversity were associated with the highest functional diversity (slope^2^ = *-*2.85 *±* 1.23, *F* = 3.27, *p* = 0.04, *R*^2^ = 0.06; Figure 3B). Other response diversity metrics were also correlated with functional (effect) diversity (Table S7).

**Figure 3:**
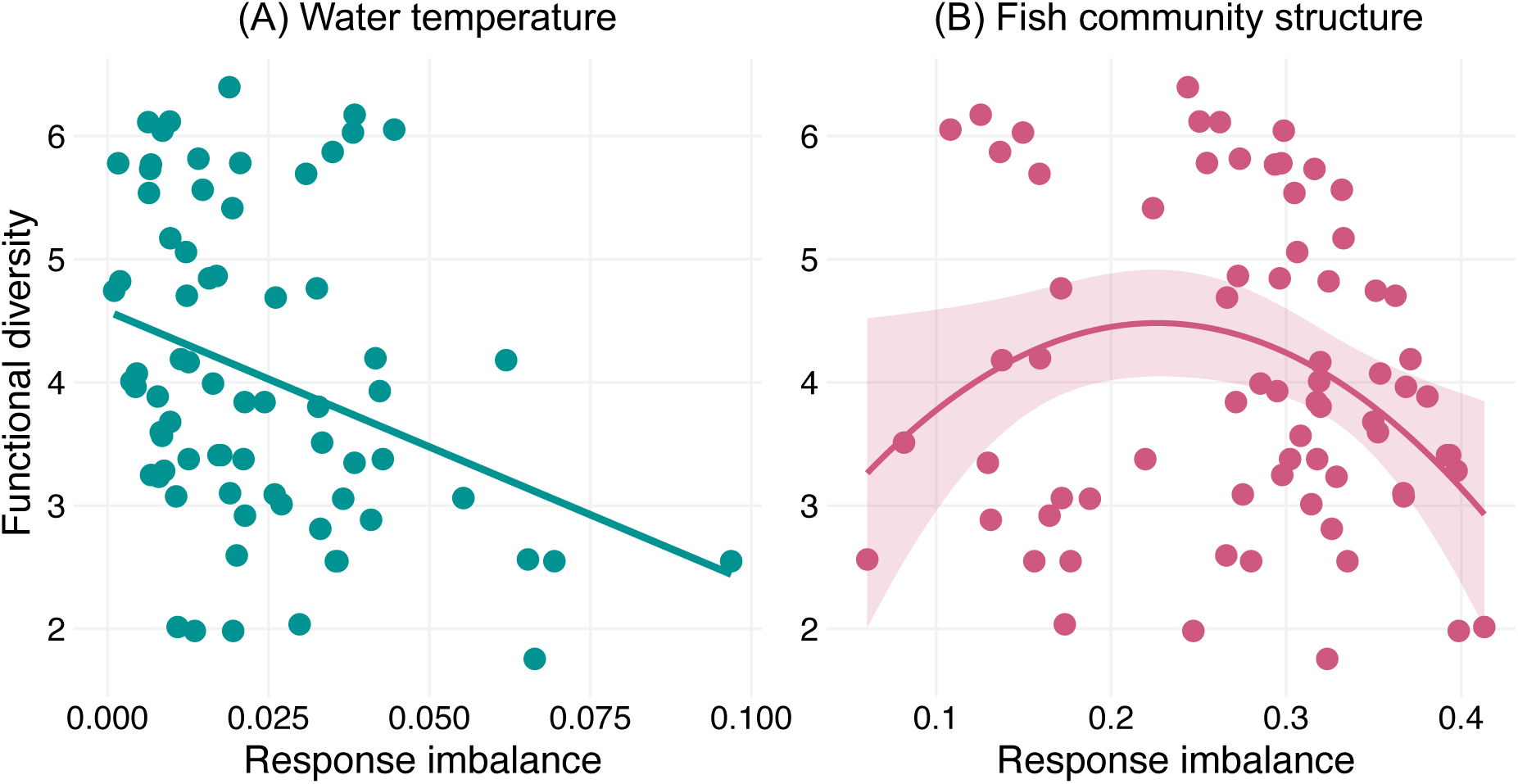
Relationship between response diversity and functional diversity. Response diversity, measured as the imbalance metric (Polazzo, Hämmig, et al. 2025), was (A) linearly negatively related to zooplankton functional diversity for water temperature (*F*_1,70_ = 8.13, *p* = 0.006, *R*^2^ = 0.09), but (B) exhibited a second order polynomial relationship with functional diversity for fish community structure (*F*_2,69_ = 3.27, *p* = 0.044, *R*^2^ = 0.06). Shading in (B) represents the 95% confidence intervals around the polynomial regression fit.

Next, we measured the relationship between species richness and functional robustness, hypothesising that response diversity provides additional explanatory power beyond that of species richness alone. We found that the response imbalance metric measured on species’ responses to water temperature was significantly negatively correlated with functional robustness in our structural equation model (path coefficient = −0.27; Figure 4A), and permutation tests to disentangle richness effects confirmed this relationship was independent of the role of species richness in shaping robustness (Figure 5A). In contrast, response imbalance under fish community structure was significantly positively related to functional robustness (path coefficient = 0.17; Figure 4B), but not independently of richness effects (Figure 5B). Species richness was consistently positively related to functional robustness across all SEMs and permutation analyses.

**Figure 4:**
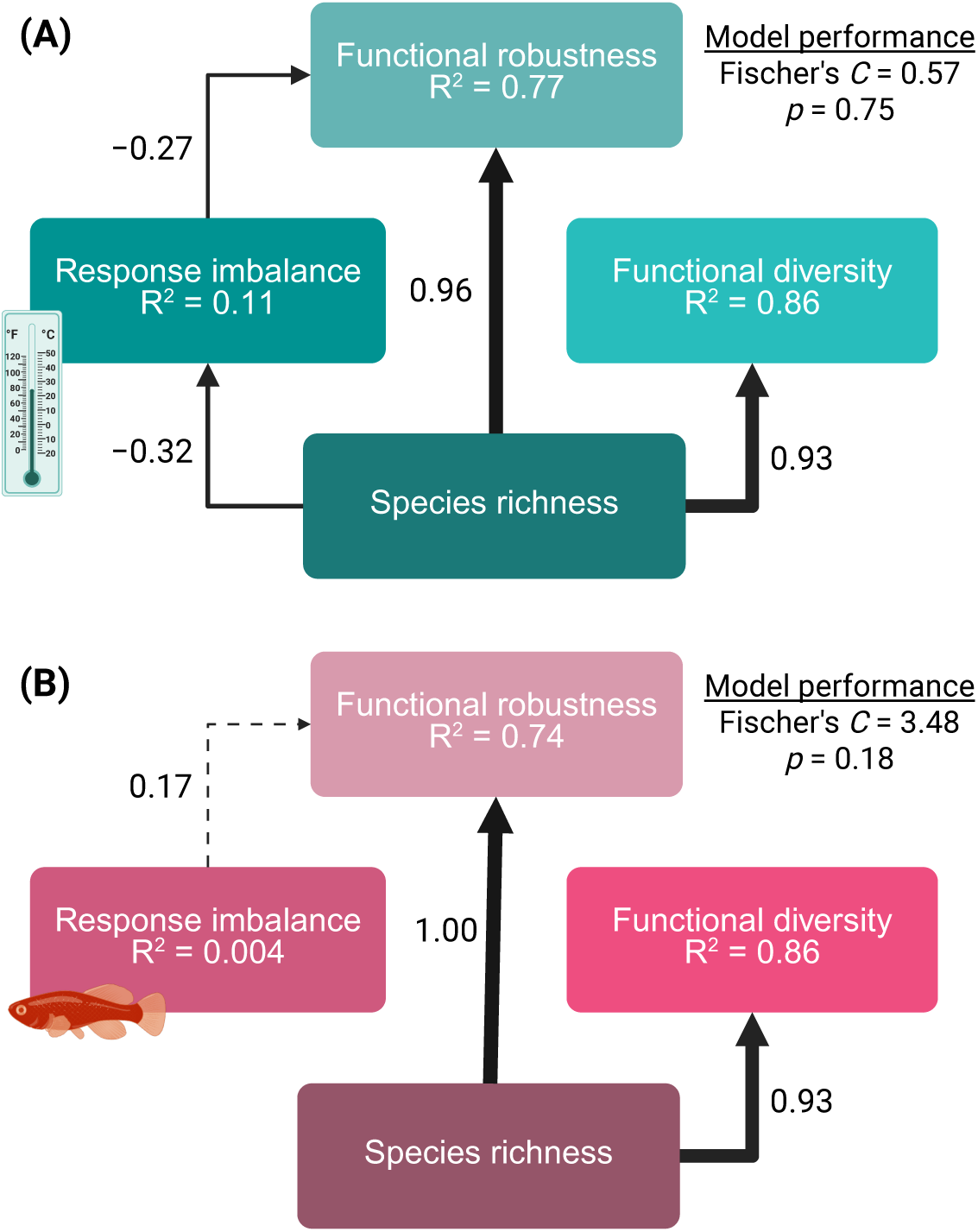
Response imbalance to an abiotic variable, but not a biotic variable, increases functional robustness. Piecewise structural equation models (SEMs) with standardised estimates of path coefficients for relationships between species richness, functional diversity, response diversity, and functional robustness. Models were fitted as OLS or GLS accounting for spatial autocorrelation (see Methods). Here, response diversity is measured as the imbalance (Polazzo, Hämmig, et al. 2025) among zooplankton species responses to (A) water temperature, and (B) the first Principal Coordinate Analysis (PCoA) axis of fish community structure. Species’ responses were estimated by joint species distribution models. Solid arrows represent significant effects in SEMs at the *p* = 0.05 threshold which remained significant after permutation tests (*i.e.* were outside the 95% confidence intervals of species randomisations). Dashed arrows represent significant effects in SEMs which fell within the bounds of the 95% confidence intervals in permutation tests. Arrow widths scale with the standardised estimate of the coefficient. *R*^2^ values indicate the explanatory power of all predictor variables on each response variable. We assessed SEM validity using Fischer’s C statistic and the *p*-value from a significance test based on the Chi-squared distribution, where *p>* 0.05 indicates model structure is appropriate (Shipley 2009). Created in BioRender.

**Figure 5:**
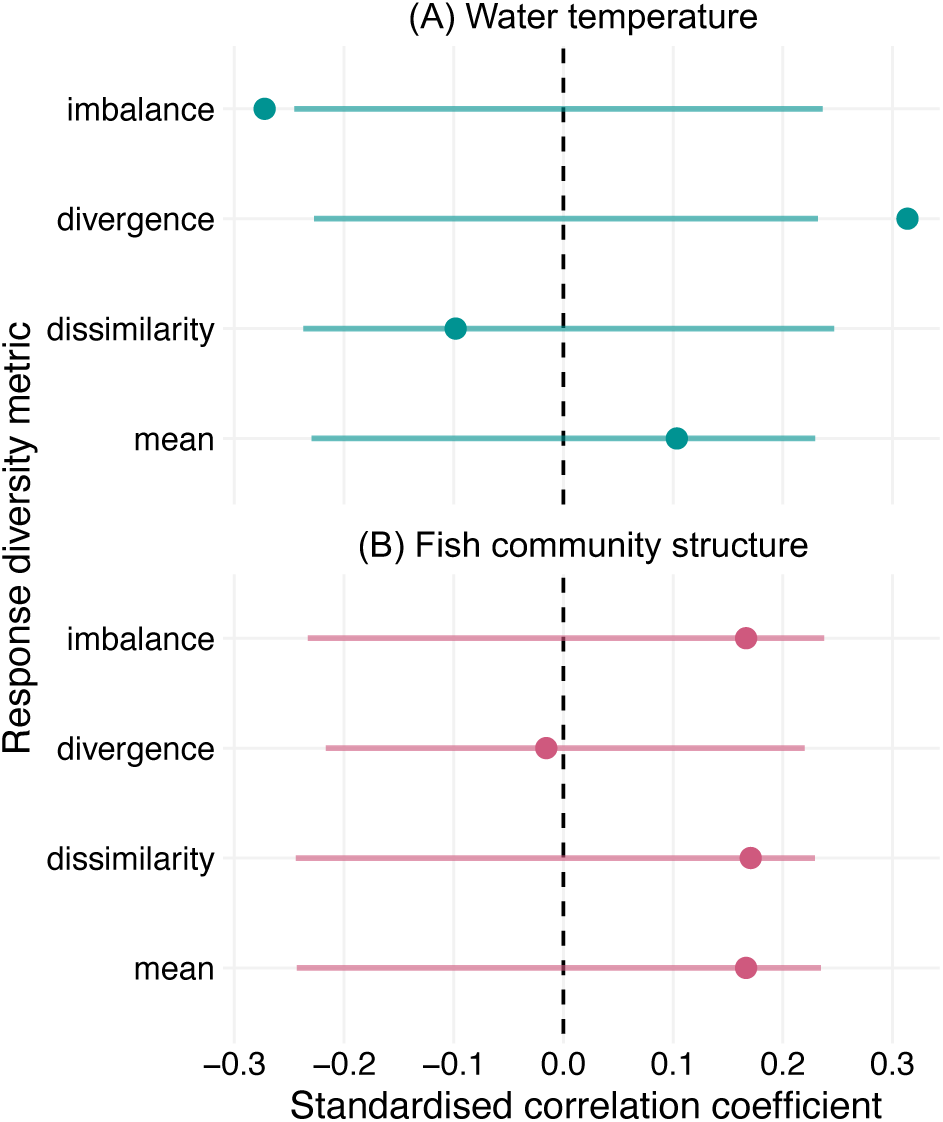
Relationships between response diversity and functional robustness compared to null models. Points represent observed standardised correlation coefficients for models between different response diversity variables and functional robustness. Grey lines indicate the 95% confidence intervals of 1,000 permutations of the response diversity-robustness relationship, where point values falling outside these lines suggest a significant effect of response diversity on functional robustness independent of the effects of species richness. Response variables were response imbalance, response divergence, response dissimilarity, and mean species response to (A) water temperature, and (B) fish community structure (PCoA axis 1). In all but two cases, species’ environmental responses and response diversity were not related to robustness independently of richness effects.

Response imbalance can be low either because species’ environmental responses are mostly weak, or alternatively because extreme negative responses are balanced by equally extreme positive responses to a given environment. These two scenarios suggest either population stability (resulting from weak environmental responses) or complementarity and asynchrony (from balanced strong responses) has a large influence in driving stability (Polazzo, Hämmig, et al. 2025). To disentangle these two possible cases for our significant response imbalance–robustness relationship (Figure 4A), we measured the range of species’ environmental responses per assemblage as the maximum ─ minimum response to water temperature. We found that low values of response imbalance are associated with larger ranges of zooplankton responses (Figure S3A), meaning a balance of positive and negative responses to water temperature underlies the stabilising effect on functional robustness here.

For other metrics of response diversity and environmental variables, we mostly found response diversity was not related to functional robustness. The one exception was response divergence to water temperature, which was significantly positively related to functional robustness (path coefficient = 0.31, Figure S4A), independently of species richness effects (Figure 5A). Response divergence to fish community structure was not related to functional robustness (Figure S4B). We found that significant SEM links between robustness and response dissimilarity (Figure S5B) and mean species response (Figure S6) were not independent of richness effects following permutation analyses (Figure 5).

We also constructed SEMs and ran permutation analyses for the environmental variables with lower relative variable importance and found that, in all but one case, response diversity metrics were not significantly related to functional robustness independently of richness effects. Response imbalance to the second PCoA axis of fish community structure was negatively related to functional robustness independently of richness effects (Figure S7), again driven by balanced positive and negative responses, rather than weak overall responses to fish community structure (Figure S3B).

## 5 Discussion

Biodiversity is widely recognised to provide insurance against environmental change and there is increasing evidence that response diversity is an important mechanism explaining this pattern. For example, communities with higher response diversity may be more resistant to pulse or press disturbance (Ross, Barros, et al. 2026; Kunze et al. 2026). However, few studies have explicitly tested the relationship between response diversity and different dimensions of ecological stability (Ross and Sasaki 2024). Functional robustness is one such dimension capturing the loss of functional diversity under a fixed perturbation strength, with consequences for the stability of ecosystem functioning (Allesina et al. 2009; Petchey and Gaston 2002a). Here, we tested whether response diversity predicts functional robustness against simulated species extinctions in 72 reservoir zooplankton assemblages across Japan, spanning a latitudinal range with broad climatic differences. In support of previous studies, we found that response diversity can provide functional robustness independently of species richness, but that this relationship is contingent on the response diversity metric used and the environmental variable considered. Craven et al. (2016) found that local response diversity and landscape connectivity maintain functional diversity under fragmentation in tree communities, partially through sampling effects—higher species richness increased response diversity by including species with distinct beneficial environmental tolerances to key anthropogenic drivers. Similarly, Altomare et al. (2021) found that savanna tree communities maintained their ecosystem functioning under fire owing to species’ diverse fire responses. Our results provide further evidence that, depending on the environmental variable examined, response diversity can predict facets of stability beyond the traditional focus on temporal stability.

Our variance partitioning approach revealed that the first dimension of fish community structure (PCoA 1) and water temperature were the most important for explaining differences in the zooplankton communities across reservoirs. Response diversity to fish community structure can arise because fish generally do not consume small zooplankton such as rotifer species and small crustaceans, or those that swim quickly, such as copepods (Zaret 1980). Fish communities structured strongly along PCoA 1—which approximates zooplankton predation pressure by omnivorous fish (Table S5)—thus will differently affect zooplankton species subjected to such fish predation versus those not. In contrast, response diversity among zooplankton in their response to water temperature arises from their physiological traits and thermal tolerances (Bonadonna et al. 2025), with low latitude (warmer) reservoirs representing a more challenging environment for cold-water specialists, for example.

In our study, the response imbalance metric—measured as the absolute community sum of derivatives of all species’ environmental responses, corrected by richness (Polazzo, Hämmig, et al. 2025)—had the strongest relationship with functional robustness. The imbalance metric captures both the direction and magnitude of species-environment responses. Accordingly, its strong correlation with stability relative to other response diversity metrics supports the idea that combining both direction and magnitude of responses best predicts stability (Ross, Petchey, et al. 2023; Polazzo, Hämmig, et al. 2025). We found response imbalance to water temperature was negatively related to functional robustness, meaning zooplankton communities with more balanced responses to water temperature were also more robust to reductions in functional diversity via species removals. Both imbalanced yet weak mean species responses (positive or negative) and balanced but stronger responses (positive *and* negative) can result in community stability, and we found that the latter was the more likely cause here (Figure S3). Determining the extent to which species’ environmental responses are balanced—producing “winners” and “losers” (Dornelas et al. 2019)—and the consequences of such balance for different facets of stability, remain critical goals in the Anthropocene.

Several metrics have been proposed to capture response diversity in terms of traits or functional responses, and also to relate response diversity to ecological stability (Ross, Petchey, et al. 2023). Yet we largely found no relationship between several response diversity metrics and functional robustness to extinctions in empirical zooplankton assemblages. Ross, Petchey, et al. (2023) found that response divergence, which captures the direction of species’ environmental responses without considering magnitude, was most clearly related to the temporal stability of aquatic ciliates, outperforming the dissimilarity metric (Leinster and Cobbold 2012), which focuses more on the magnitude of differences in species responses. In support of this, we found that zooplankton assemblages with more divergent responses to water temperature were more functionally robust, whereas the magnitude of such differences (measured by response dissimilarity) tended to be unrelated to robustness. This result also supports our above finding that a balance of positive and negative species responses to water temperature can confer functional robustness. As with response dissimilarity, the mean species environmental response was not related to functional robustness in our study, despite its inferred importance for shaping stability and coexistence in other studies (De Laender et al. 2023; Kunze et al. 2026).

In this study, the choice of environmental variable largely did not change our results—besides water temperature, response diversity to each environmental axis was consistently not correlated with functional robustness independently of richness (Figure S6). Although we chose predictor variables that we hypothesized to explain zooplankton species’ distributions and their responses to the environment, we may have missed important variables that can better explain the functional robustness of these zooplankton assemblages. However, response diversity was significantly related to functional (effect) diversity in our study, indicating that species’ environmental responses should propagate to stabilise functions (De Bello et al. 2021). The lack of consistent response diversity effects on functional robustness after accounting for richness, despite the relationship of response diversity with functional diversity, suggests that response–effect trait linkages may not scale uniformly to influence different dimensions of stability (Suding et al. 2008; Mori et al. 2013; De Bello et al. 2021). Along most environmental axes considered here, these findings collectively suggest that response diversity may contribute more strongly to shaping functional structure than to conferring functional robustness against species losses.

Though parameterised on empirical data, our analysis framework may have failed to realistically capture the functional robustness of real-world zooplankton assemblages. We simulated random species removals on zooplankton assemblages and measured functional robustness based on the loss of functional diversity under these extinctions. Yet, non-random extinctions are common in nature, and extinction effects propagate further under realistic extinction sequences where species are lost based on their trophic links, relative abundance, or functional traits (Dunne et al. 2002; Berg et al. 2015). Recently, Hsieh et al. (2026) found response diversity of phytoplankton and zooplankton communities stabilised biomass, but this effect was contingent on time-varying environmental conditions. Our modelling approach used time-invariant environmental parameters, missing possible temporal effects of fluctuating environmental conditions on robustness. Similarly, realistic scenarios considering multiple simultaneous environmental changes may reveal the joint effects of several axes of species’ environmental responses on stability (Polazzo, Limberger, et al. 2024).

Another explanation for the inconsistent relationships between response diversity and functional robustness in our study may be that we did not recover realistic species-environment responses from our correlative species distribution models. Modelled responses of probability of species occurrence to the environment, representing estimates of realized niches, do not directly capture mechanistic physiological or functional responses. Additionally, our estimated species responses may be biased toward or against certain environmental values due to co-occurrence patterns with heterospecifics (Pollock et al. 2014) or the realised distribution of conditions in environmental parameter space (*i.e.* the “realised environment”, Guisan and Thuiller 2005). Mechanistic response curves based on mean individual performance or biomass, for example, should provide better estimates of fundamental niches and thus better predict functional stability than indirect estimates (Ross, Petchey, et al. 2023). For example, experimentally measuring species’ functional responses can mechanistically link response diversity and stability (Leary and Petchey 2009; Polazzo, Hämmig, et al. 2025), but experiments may overestimate environmental change impacts through unrealistic study designs (Korell et al. 2020; but see Collins et al. 2022). In contrast, our environmental variables were parameterised with data from biodiversity monitoring (see *e.g.*, Suzuki, Ichiyanagi, et al. 2025), affording us insight into the environmental conditions that structure wild zooplankton assemblages. Mechanistic response data for zooplankton species were unavailable here. As this is commonly the case in other systems, the workflow we use here, based on species distribution models, offers a viable alternative for inference.

In our study, the relationship between response diversity and robustness depended both on the choice of metric and the environmental axis considered. Given that response diversity only rarely drove functional robustness, and also that species richness is not the direct mechanism by which biodiversity confers stability (McCann 2000), additional explanatory variables are needed to disentangle species richness and stability in our study. We used presence-absence data and measured response diversity from the environmental parameters that predicted species occurrence. In this way, we could not determine any effects of species abundance distributions, such as the known influence of dominant species dynamics on stability (Sasaki and Lauenroth 2011; Luo et al. 2025). Recently, Genung and Winfree (2025) found that a single, highly dominant bee species with high abundance made a disproportionate contribution to response diversity that stabilised crop pollination, demonstrating that abundance patterns are also crucial to consider when making predictions of stability based on response diversity. Functional robustness can also be considered in a network context (Allesina et al. 2009; Domínguez-García et al. 2019; Ross, Arnoldi, et al. 2021), allowing consideration of response diversity as a stability driver in ecological networks with diverse interspecific interactions (Danet et al. 2025), though we could not take this approach here due to lack of network data. Including such facets of zooplankton assemblage structure would give more power to conclusively accept or reject the hypothesis that response diversity drives robustness in these reservoirs.

## 6 Conclusion

Response diversity has been proposed as a causal mechanism explaining the stability of empirical ecological communities. Owing to sampling effects, species-rich communities have a higher probability of including species with diverse environmental responses, as well as those with diverse functional roles (Mori et al. 2013). If species’ environmental responses and their functional roles are correlated, as we found here, response diversity can confer stability to environmental change (De Bello et al. 2021). However, we found only limited empirical evidence that response diversity relates to the functional robustness of zooplankton assemblages in reservoirs across the Japanese archipelago. In contrast to work showing positive effects of response diversity on resilience and functional persistence (Craven et al. 2016; Altomare et al. 2021), and also on temporal stability (*e.g.*, Leary and Petchey 2009; Sasaki, Lu, et al. 2019; White et al. 2023; Genung and Winfree 2025), our study suggests that response diversity can confer functional robustness to species extinctions independently of species richness effects, but perhaps not widely and only for certain environmental axes. This study highlights the importance of evaluating the predictive capacity of response diversity as a driver of ecological stability beyond temporal invariability, and also of testing response diversity metrics derived from diverse sources of species-environment relationships.

## Author Contributions

**S.R.P-J. Ross:** Conceptualisation, Methodology, Software, Formal analysis, Investigation, Resources, Writing - Original Draft, Writing - Review & Editing, Visualisation, Project administration, Funding acquisition. **H. Suzuki:** Methodology, Data Curation, Writing-Original Draft, Writing - Review & Editing, Visualisation. **J. Urabe:** Methodology, Data Curation, Writing - Original Draft, Writing - Review & Editing, Supervision. **J.M. Kass:** Conceptualisation, Methodology, Software, Validation, Formal analysis, Investigation, Resources, Writing - Original Draft, Writing - Review & Editing, Visualization, Funding acquisition.

## Acknowledgments

We would like to thank Tomonori Osugi, Hidetaka Ichiyanagi, and the staff of the Water Resources Environment Center for creating and managing the River Environmental Database used in this study. We thank Laura Antaõ, Shaopeng Wang, and Francesco Polazzo for helpful discussion. This study was supported by a SHINKA research collaboration grant and a SHINKA 2nd top-up grant awarded to SRPJR and JMK by the Okinawa Institute of Science and Technology Graduate University (OIST) and Tohoku University, and a Dam-research fund by the Water Resources Environment Center, Japan, to JU, with additional partial support from a BES Large Grant (LRB22/1007) to SRPJR and the Environment Research and Technology Development Fund (JPMEERF20214003) to JU. SRPJR was supported by subsidy funding to OIST. This paper in part results from the activities and support of the Response Diversity Network (https://responsediversitynetwork.github.io/RDN-website/).

## Data and Code Availability

All the data used in this study were obtained from disclosed sources including the Database of Dams in Japan (http://mudam.nilim.go.jp/home) for the location and watershed areas; the Japan Dam Foundation (http://damnet.or.jp/) for the dam specification data; and the River Environmental Database (http://www.nilim.go.jp/lab/fbg/ksnkankyo/ index.html) for water chemistry, fish, and zooplankton data. All code needed to reproduce the analyses in this study are open and available via Zenodo: (link will be added upon acceptance). Trait data sources are listed in the Supporting Information. Sources for environmental parameters are listed in the Methods section.

