## Supplementary material for "Response diversity inconsistently predicts functional robustness to simulated zooplankton extinctions": _

Table S1: Summary of geographical position, dam characteristics, trophic conditions (TN: total nitrogen, TP: total phosphorus, and chlorophyll $a$), and water temperature of the 73 reservoirs examined.

| Environmental variable | Mean value | Median value | Value range |
|---|---|---|---|
| Latitude | - | 35.866 | 26.48−44.114 |
| Longitude | - | 137.881 | 127.952−143.387 |
| Altitude (m) | 329 | 285 | 28−976 |
| Reservoir volume ($\times 10^3$ $m^3$) | 59,672 | 42,550 | 4,000−289,000 |
| Dam height (m) | 88 | 85 | 24−158 |
| Dam width (m) | 334 | 310 | 128−1,480 |
| TN (mg/L) | 0.469 | 0.383 | 0.142−1.553 |
| TP (mg/L) | 0.016 | 0.013 | 0.003−0.09 |
| Chlorophyll-$a$ ($\mu$g/L) | 4.1 | 2.5 | 0.08−35.43 |
| Water temperature ($^o$C) | 17.1 | 17.3 | 8.9−24 |

Table S2: **Zooplankton functional trait data included in this study**. We selected traits that related to species' functional roles (*i.e.*, functional effect traits) rather than species' environmental responses (Suding et al. 2008; Oliver et al. 2015).

| Trait | Value range | Justification | Source |
|---|---|---|---|
| Mean body size | 90 $\mu$m – 1400 $\mu$m | Body size affects metabolic rate (Lampert 1984) and feeding rate (Peters and Downing 1984). | Edmondson (1959), Thienemann (1974), Mizuno and Takahashi (2000), Tanaka and Makita (2017), and Tanaka (2022). |
| Food preference | First two principal component analysis (PCA) axes based on percentage diet composition data from blood-feeding, carnivory, detritivory, herbivory, omnivory | Indicates trophic level and feeding interactions for each taxon. | Salt et al. (1978) and Suzuki et al. (2025). |
| Feeding strategy | Filter, Raptorial, Browsing | Affects food availability and feeding efficiency (Kiørboe 2011). | Suzuki et al. (2025). |
| Swimming mode | Darting, Hopping, Drifting | Affects food accessibility and feeding efficiency (Kiørboe 2011) and is linked to metabolic rate. | Suzuki et al. (2025). |

Table S3: Results of the principal component analysis of the dam structural characteristics for Japanese reservoirs, showing standard deviation, variance explained, and scores for each dimension.

| | Axis 1 | Axis 2 | Axis 3 | Axis 4 |
|---|---|---|---|---|
| Standard deviation | 1.28 | 1.047 | 0.94 | 0.62 |
| Variance explained | 40.7% | 27.4% | 22.2% | 9.7% |
| Watershed area | 0.004 | 0.90 | -0.27 | 0.34 |
| Dam volume | -0.67 | 0.19 | -0.25 | -0.67 |
| Dam height | -0.63 | -0.32 | -0.28 | 0.64 |
| Dam width | -0.38 | 0.22 | 0.89 | 0.12 |

Table S4: Results of the principal component analysis of the water nutrient variables for Japanese reservoirs, showing standard deviation, variance explained, and scores for each dimension.

| | Axis 1 | Axis 2 | Axis 3 |
|---|---|---|---|
| Standard deviation | 1.37 | 0.90 | 0.56 |
| Variance explained | 62.5% | 27.1% | 10.3% |
| TN | 0.65 | -0.23 | -0.72 |
| TP | 0.63 | -0.37 | 0.68 |
| Chlorophyll-*a* | 0.43 | 0.90 | 0.10 |

Table S5: Results of the first three dimensions of the principal coordinate analysis of the fish community in Japanese reservoirs, showing eigenvalues, variance explained and scores for each dimension.

| | Axis 1 | Axis 2 | Axis 3 |
|---|---|---|---|
| Eigenvalue | 14.3 | 7 | 5.5 |
| Variance explained | 21.7% | 10.4% | 8.1% |
| *Phoxinus* sp. | 0.059 | 0.191 | 0.307 |
| *Plecoglossus altivelis* | 0.274 | 0.235 | -0.101 |
| *Salvelinus* sp. | -0.294 | 0.194 | 0.161 |
| *Gymnogobius* sp. | 0.120 | 0.060 | -0.059 |
| *Leuciscus hakonensis/L. ezoe* | -0.055 | 0.275 | 0.229 |
| *Anguilla japonica* | 0.099 | 0.044 | -0.051 |
| *Zacco platypus* | 0.287 | 0.304 | 0.076 |
| *Micropterus salmoides* | 0.250 | 0.201 | -0.076 |
| *Cottus pollux* | -0.084 | -0.036 | 0.183 |
| *Pseudogobio ecocinus* | 0.258 | 0.183 | -0.003 |
| *Nipponocypris temminckii/N. sieboldii* | 0.253 | 0.162 | -0.122 |
| *Pseudobagrus fulvidraco* | 0.171 | 0.160 | -0.116 |
| *Carassius cuvieri* | 0.145 | 0.201 | 0.151 |
| *Carassius* sp. | 0.158 | -0.169 | 0.175 |
| *Cyprinus carpio* | 0.112 | -0.043 | 0.320 |
| *Salmo masou masou* | -0.053 | 0.135 | 0.309 |
| *Salmo masou macrostomus* | 0.040 | 0.059 | 0.007 |
| *Cobitis biwae* | 0.059 | 0.081 | 0.089 |
| *Squalidus* sp. | 0.246 | 0.193 | -0.103 |
| *Biwia zezera* | 0.059 | 0.094 | 0.086 |
| *Gnathopogon* sp. | 0.091 | 0.157 | 0.212 |
| *Misgurnus anguillicaudatus* | -0.020 | 0.106 | 0.387 |
| *Odontobutis obscura obscura* | 0.096 | 0.065 | -0.042 |
| *Silurus asotus* | 0.223 | 0.174 | -0.053 |
| *Hemibarbus labeo barbus* | 0.182 | 0.212 | 0.053 |
| *Oncorhynchus mykiss* | -0.163 | 0.290 | -0.036 |
| *Tridentiger kuroiwae brevispinis* | 0.194 | 0.203 | -0.026 |
| *Opsariichthys uncirostris* | 0.236 | 0.213 | -0.085 |
| *Noemacheilus barbatulus toni* | -0.132 | 0.078 | -0.099 |
| *Lepomis macrochirus* | 0.212 | 0.155 | -0.148 |
| *Pungtungia herzi* | 0.160 | 0.108 | -0.097 |
| *Oryzias latipes* | 0.051 | -0.042 | -0.010 |
| *Pseudorasbora parva* | 0.051 | 0.114 | 0.285 |
| *Rhinogobius* sp. | 0.224 | -0.236 | 0.086 |
| *Hypomesus nipponensis* | -0.050 | 0.163 | 0.346 |

Table S6: Site loadings for the first three dimensions of the principal coordinate analysis of the fish community in Japanese reservoirs, showing scores for each dam along each dimension.

| | Axis 1 | Axis 2 | Axis 3 |
|---|---|---|---|
| Aha | -0.13 | -0.92 | 0.00 |
| Arakawa | -0.15 | -0.81 | -0.37 |
| Aseishigawa | -0.34 | -0.14 | 0.39 |
| Benoki | -0.13 | -0.92 | 0.00 |
| Fujiwara | -0.37 | -0.12 | 0.50 |
| Fukuichi | -0.11 | -0.88 | -0.23 |
| Gassan | -0.41 | -0.35 | 0.15 |
| Gosyo | -0.005 | 0.08 | 0.39 |
| Haji | 0.50 | 0.20 | 0.017 |
| Hanezi | -0.13 | -0.92 | 0.00 |
| Hattabara | 0.53 | 0.06 | -0.25 |
| Hinachi | 0.52 | 0.06 | -0.45 |
| Hitokura | 0.45 | 0.13 | -0.37 |
| Hiyoshi | 0.52 | 0.10 | -0.12 |
| Houheikyo | -0.86 | 0.29 | -0.66 |
| Ikari | 0.09 | 0.30 | 0.14 |
| Ikeda | 0.43 | 0.10 | -0.05 |
| Iwaonai | -0.85 | 0.37 | -0.28 |
| Iwaya | 0.43 | 0.13 | 0.06 |
| Izarigawa | -0.88 | 0.35 | -0.49 |
| Jyozankei | -0.41 | -0.24 | 0.19 |
| Kamafusa | -0.06 | 0.17 | 0.37 |
| Kanayama | -0.47 | 0.04 | 0.25 |
| Kanna | -0.14 | -0.96 | -0.23 |
| Kanoko | -0.70 | 0.02 | -0.05 |
| Kawaji | -0.08 | 0.19 | 0.37 |
| Kawamata | -0.09 | 0.13 | 0.23 |
| Koshibu | -0.13 | 0.20 | 0.04 |
| Managawa | -0.12 | 0.25 | -0.14 |
| Matsubara | 0.47 | 0.09 | -0.11 |
| Midorikawa | 0.39 | 0.07 | 0.03 |
| Miharu | 0.19 | 0.05 | 0.22 |
| Miwa | -0.047 | 0.14 | 0.20 |
| Miyagase | 0.016 | 0.16 | 0.12 |
| Muroo | 0.54 | 0.09 | -0.05 |
| Nagashima | -0.26 | 0.008 | 0.18 |
| Naramata | -0.22 | 0.13 | 0.53 |

| | Axis 1 | Axis 2 | Axis 3 |
|---|---|---|---|
| Narugo | 0.008 | 0.35 | -0.02 |
| Nukui | 0.35 | 0.056 | 0.054 |
| Nunome | 0.35 | 0.15 | 0.054 |
| Oodo | 0.45 | 0.06 | -0.13 |
| Ooishi | -0.51 | -0.16 | 0.17 |
| Ookawa | 0.28 | 0.12 | 0.25 |
| Pirika | -0.82 | 0.38 | -0.11 |
| Sagurigawa | -0.64 | 0.22 | 0.27 |
| Sameura | 0.38 | 0.032 | -0.10 |
| Sarudani | 0.40 | 0.10 | -0.039 |
| Satsunaigawa | -0.79 | 0.26 | -0.62 |
| Shijushida | 0.14 | -0.13 | 0.18 |
| Shimokubo | 0.28 | 0.22 | 0.15 |
| Shimouke | 0.42 | 0.088 | -0.025 |
| Shinguu | 0.46 | 0.024 | -0.24 |
| Shintoyone | 0.33 | 0.055 | 0.037 |
| Sonohara | -0.023 | 0.098 | 0.45 |
| Sugesawa | 0.06 | -0.078 | 0.09 |
| Surikamigawa | -0.24 | 0.058 | 0.37 |
| Takayama | 0.60 | 0.069 | -0.22 |
| Takisato | -0.44 | -0.041 | 0.044 |
| Tase | 0.20 | 0.12 | 0.29 |
| Tedorigawa | -0.31 | 0.047 | 0.35 |
| Terauchi | 0.40 | 0.089 | 0.023 |
| Tokachi | -0.88 | 0.35 | -0.49 |
| Tomada | 0.43 | 0.007 | -0.14 |
| Tomisato | 0.12 | -0.22 | 0.0018 |
| Tsuruta | 0.55 | -0.0012 | -0.20 |
| Yabakei | 0.46 | 0.010 | -0.39 |
| Yagisawa | -0.20 | -0.030 | 0.42 |
| Yanase | 0.37 | 0.030 | -0.25 |
| Yasaka | 0.51 | 0.053 | -0.25 |
| Yokokawa | -0.73 | 0.21 | -0.33 |
| Yokoyama | 0.38 | 0.014 | -0.039 |
| Yuda | -0.32 | -0.28 | -0.13 |

Table S7: **Relationships between functional diversity and response diversity for different metrics and environmental variables**. Functional diversity was modelled as a functional of response variables response imbalance (Polazzo et al. 2025), response dissimilarity (Leinster and Cobbold 2012), response divergence (Ross et al. 2023), and mean species response. We compared linear models with those including a 2nd order polynomial term, and evaluated model fit with $F$-tests and Akaike's Information Criterion (AIC). Polynomial models that performed significantly better ($p$ <0.05, $\delta$AIC < 2) are marked here with "$^2$". $F$-tests and $p$-values with significance thresholds (* <0.05, ** <0.01, *** <0.001) for full models are presented here with adjusted $R^2$ values indicating goodness-of-fit.

| Variable(s) | $F$ | $p$-val. (sig.) | adj.$R^2$ |
|---|---|---|---|
| Imbalance (water temp.) | 8.13 | 0.006 (**) | 0.09 |
| Divergence$^2$ (water temp.) | 8.18 | 0.032 (*) | 0.17 |
| Dissimilarity (water temp.) | 5.23 | 0.025 (*) | 0.06 |
| Mean response$^2$ (water temp.) | 5.64 | 0.002 (**) | 0.12 |
| Imbalance$^2$ (fish PCoA1) | 3.27 | 0.024 (*) | 0.06 |
| Divergence (fish PCoA1) | 0.58 | 0.45 (N.S.) | 0.01 |
| Dissimilarity (fish PCoA1) | 4.88 | 0.031 (*) | 0.05 |
| Mean response$^2$ (fish PCoA1) | 3.27 | 0.044 (*) | 0.06 |

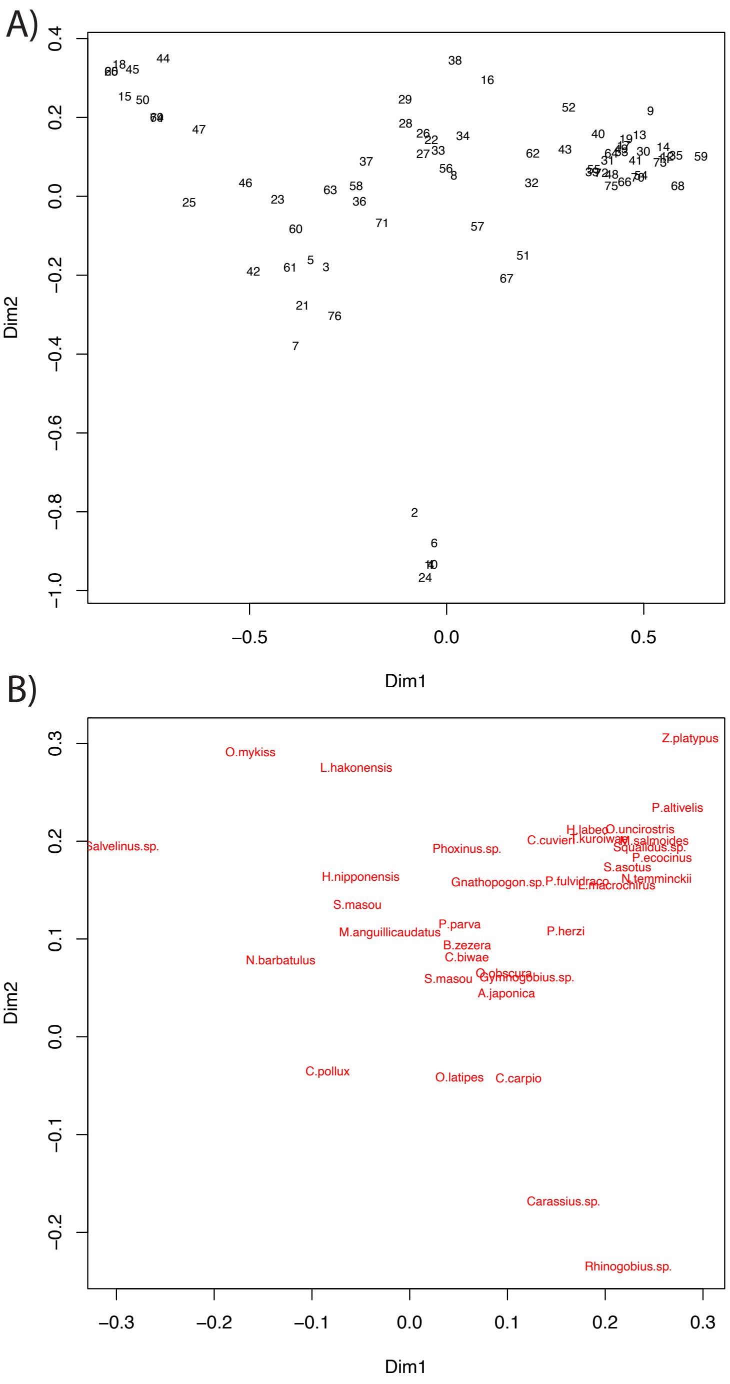


Figure S1: **Fish community structure ordination**. First two axes of a principal coordinate analysis (PCoA) of fish community structure, with loads for different sites (A) and species (B) on the first two PCoA axes. See Tables S6 and S6, respectively, for scores on each axis.

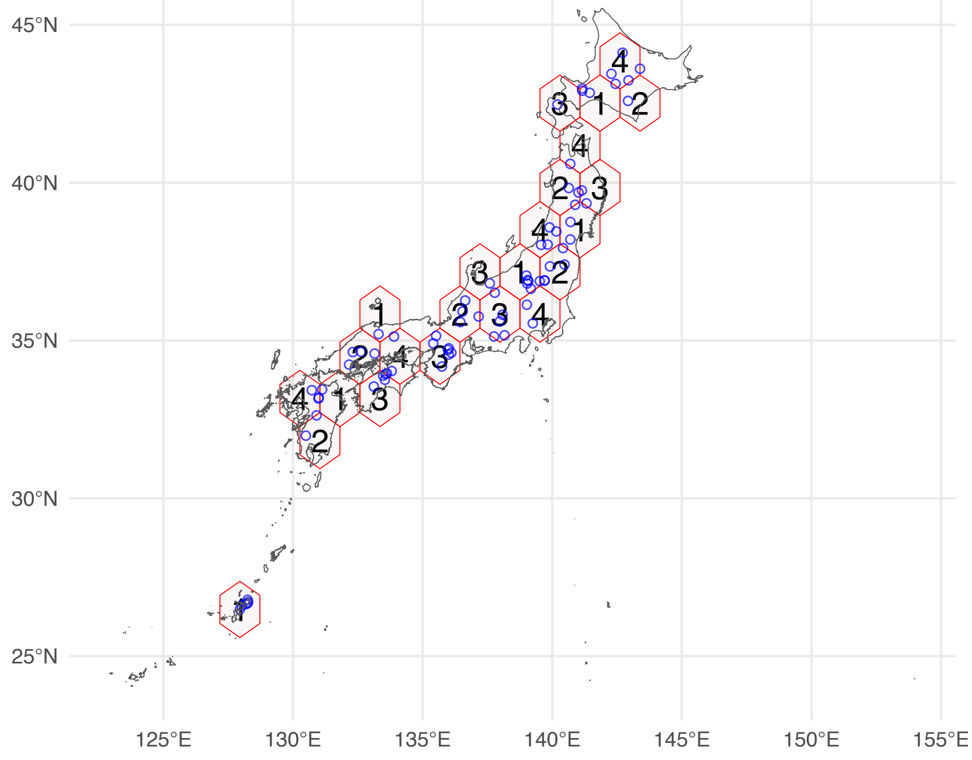


Figure S2: **Spatial partitions used for cross-validation to evaluate the joint species distribution models**. Numbers refer to different partitions. This ensures that evaluations are spatially distinct, thus reducing spatial dependencies between validation points, and ultimately results in a more difficult prediction exercise for the model. Reservoir sites records are shown with blue points.

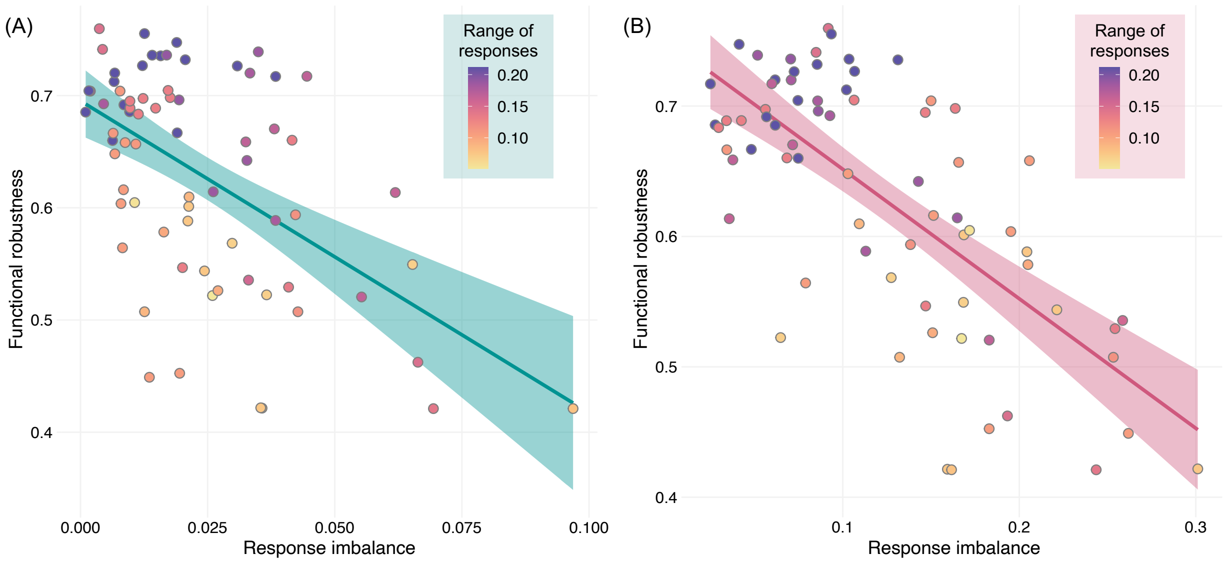


Figure S3: **Significant relationships between functional robustness and response imbalance**. Response imbalance calculated on zooplankton responses to (A) water temperature, and (B) fish community structure (PCoA axis 2) were significantly related to functional robustness independently of richness (see Figures 5 and S7). Shading represents the 95% confidence intervals around the regression fit. Point colours indicate the range of responses (maximum – minimum) to water temperature among the species in each reservoir community, where darker colours a larger range of responses. Response imbalance can be low either because responses are balanced between positives and negatives, or because responses are overall weak (Polazzo et al. 2025). Here, larger ranges of responses were concentrated at low imbalance (and high robustness), suggesting that response imbalance is related to robustness through a balance of positive and negative responses, rather than through weak overall assemblage responses.

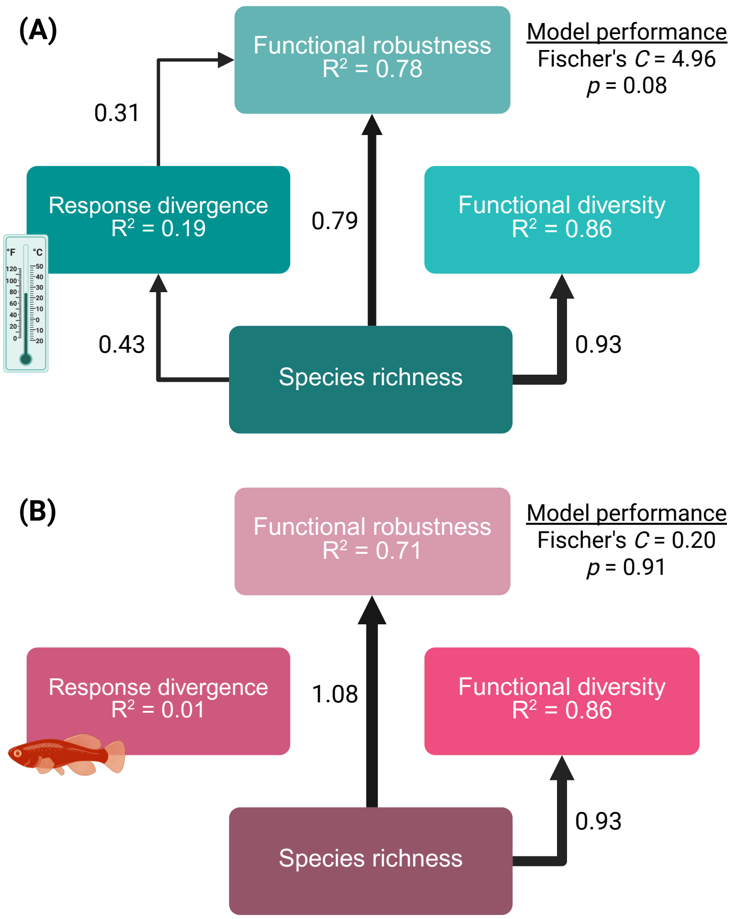


Figure S4: **Response divergence to water temperature, but not fish community structure, increases functional robustness**. Piecewise structural equation models (SEMs) with standardised estimates of path coefficients for relationships between species richness, functional diversity, response diversity, and functional robustness. Models were fitted as OLS or GLS accounting for spatial autocorrelation (see Methods). Here, response diversity is measured as the divergence (Ross et al. 2023) among zooplankton species responses to (A) water temperature, and (B) the first Principal Coordinate Analysis (PCoA) axis of fish community structure. Species' responses were estimated by joint species distribution models. Solid arrows represent significant effects in SEMs at the $p = 0.05$ threshold, which remained significant after permutation tests (*i.e.* were outside the 95% confidence intervals of species randomisations). Dashed arrows represent significant effects in SEMs which fell within the bounds of the 95% confidence intervals in permutation tests. Arrow widths scale with the standardised estimate of the coefficient. $R^2$ values indicate the explanatory power of all predictor variables on each response variable. We assessed SEM validity using Fischer's C statistic and the p-value from a significance test based on the Chi-squared distribution, where $p > 0.05$ indicates model structure is appropriate (Shipley 2009). Created in BioRender.

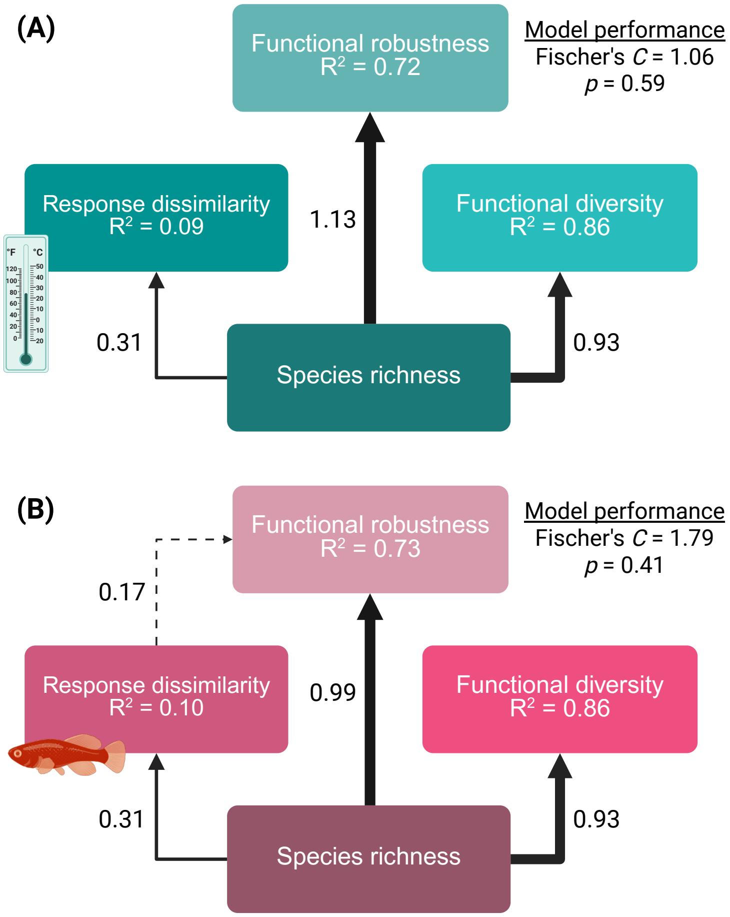


Figure S5: **Response dissimilarity does not increase functional robustness independently of richness**. Piecewise structural equation models (SEMs) with standardised estimates of path coefficients for relationships between species richness, functional diversity, response diversity, and functional robustness. Models were fitted as OLS or GLS accounting for spatial autocorrelation (see Methods). Here, response diversity is measured as the richness-only similarity metric (Leinster and Cobbold 2012) adapted for the study of response diversity (Ross et al. 2023) among zooplankton species responses to (A) water temperature, and (B) the first Principal Coordinate Analysis (PCoA) axis of fish community structure. Species' responses were estimated by joint species distribution models. Solid arrows represent significant effects in SEMs at the $p = 0.05$ threshold, which remained significant after permutation tests (*i.e.* were outside the 95% confidence intervals of species randomisations). Dashed arrows represent significant effects in SEMs which fell within the bounds of the 95% confidence intervals in permutation tests. Arrow widths scale with the standardised estimate of the coefficient. $R^2$ values indicate the explanatory power of all predictor variables on each response variable. We assessed SEM validity using Fischer's C statistic and the p-value from a significance test based on the Chi-squared distribution, where $p > 0.05$ indicates model structure is appropriate (Shipley 2009). Created in BioRender.

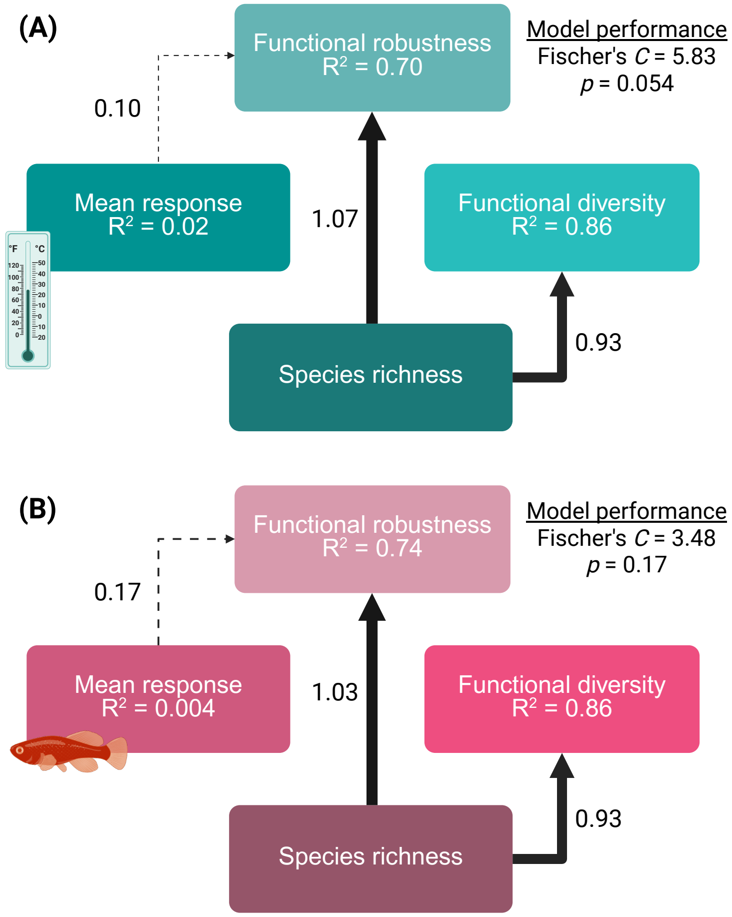


Figure S6: **Mean species response does not increase functional robustness independently of richness**. Piecewise structural equation models (SEMs) with standardised estimates of path coefficients for relationships between species richness, functional diversity, mean response, and functional robustness. Models were fitted as OLS or GLS accounting for spatial autocorrelation (see Methods). Here, mean response is the mean of species' responses in a zooplankton assemblage (Kunze et al. 2026) to (A) water temperature, and (B) the first Principal Coordinate Analysis (PCoA) axis of fish community structure. Species' responses were estimated by joint species distribution models. Solid arrows represent significant effects in SEMs at the $p = 0.05$ threshold, which remained significant after permutation tests (*i.e.* were outside the 95% confidence intervals of species randomisations). Dashed arrows represent significant effects in SEMs which fell within the bounds of the 95% confidence intervals in permutation tests. Arrow widths scale with the standardised estimate of the coefficient. $R^2$ values indicate the explanatory power of all predictor variables on each response variable. We assessed SEM validity using Fischer's C statistic and the p-value from a significance test based on the Chi-squared distribution, where $p > 0.05$ indicates model structure is appropriate (Shipley 2009). Created in BioRender.

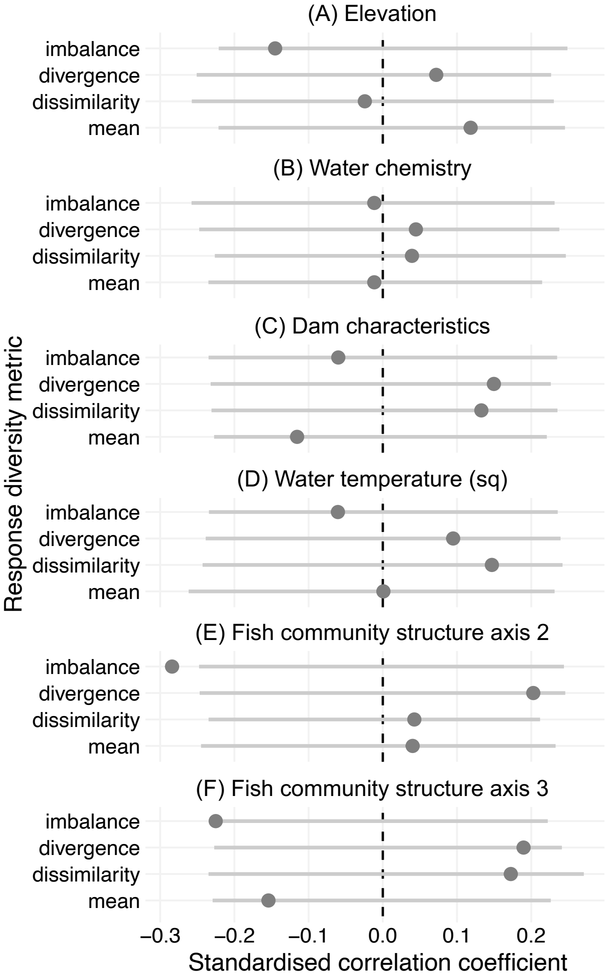


Figure S7: **Relationships between response diversity and functional robustness compared to null models**. Points represent observed standardised correlation coefficients for models between different response diversity variables and functional robustness. Grey lines indicate the 95% confidence intervals of 1,000 permutations of the response diversity-robustness relationship, where point values falling outside these lines suggest a significant effect of response diversity on functional robustness independent of the effects of species richness. Response variables were response imbalance, response divergence, response dissimilarity, and mean species response to (A) elevation, (B) water chemistry, (C) dam/reservoir characteristics, (D) the second order polynomial term for water temperature, (E) the second PCoA axis for fish community structure, and (F) the third PCoA axis for fish community structure. In almost all cases, species' environmental responses and response diversity were not related to robustness independently of richness effects. Response imbalance—measured on PCoA axis 2 of the fish community structure—was significantly related to functional robustness independently of richness; lower imbalance values (more balanced responses) promoted robustness.
